# Tissue-type specific accumulation of the plastoglobular proteome, transcriptional networks and plastoglobular functions

**DOI:** 10.1101/2021.02.05.430006

**Authors:** Elena J.S. Michel, Lalit Ponnala, Klaas J. van Wijk

**Affiliations:** School of Integrative Plant Sciences (SIPS), Section of Plant Biology, Cornell University, Ithaca, New York 14853, USA; Viqstra, Inc., Staten Island, NY 10304

**Keywords:** plastoglobules, co-expression networks, isoprenoids, chloroplast, proteome, senescence

## Abstract

Plastoglobules (PGs) are dynamic protein-lipid micro-compartments in plastids enriched for isoprenoid-derived metabolites. Chloroplast PGs support formation, remodeling and controlled dismantling of thylakoids during developmental transitions and environmental responses. However, the specific molecular functions of most PG proteins are still poorly understood. This study harnesses recent co-mRNA expression from ATTED-II using combined microarray and RNAseq information on an updated inventory of 34 PG proteins, as well as proteomics data across 30 Arabidopsis tissue types from ATHENA. Hierarchical clustering based on relative abundance for the PG proteins across non-photosynthetic and photosynthetic tissue types showed their coordinated protein accumulation across Arabidopsis parts, tissue types, development and senescence. We generated multiple mRNA-based networks by applying different coefficient thresholds; functional enrichment was determined for each network and PG gene. Combined analysis of these stringency networks identified a central hub and four peripheral modules. Enrichment of specific nuclear transcription factors (*e*.*g*. Golden2-like) and support for cross-talk between PGs and the plastid gene expression was observed, and specific ABC1 kinases seem part of a light signaling network. Examples of other specific findings are that FBN7b is involved with upstream steps of tetrapyrrole biosynthesis and that ABC1K9 is involved in starch metabolism.

**Highlight:** The plastoglobular proteome shows coherent tissue-specific accumulation, whereas combined analysis of transcriptional co-expression networks, at different stringencies and following in-depth functional annotation, associate selected plastoglobular proteins to specific metabolic functions.

## Introduction

Plastoglobules (PGs) are dynamic protein monolayer lipid micro-compartments in all plastid types, reviewed in (van Wijk and Kessler, 2017). In chloroplasts, they swell from the outer thylakoid membrane leaflet and respond dynamically to abiotic and developmental changes. PGs have a distinct proteome of more than 30 proteins including tocopherol cyclase (VTE1), carotenoid cleavage 4 (CCD4), NADPH oxidoreductase (NDC1), six members of the atypical ‘Activity of BC1’ kinase family (ABC1K1,3,5,6,7,9) and multiple proteins with unknown function (van Wijk and Kessler, 2017). The ancestral role of the ABC1K is likely regulation of quinone synthesis, such as naphthoquinones in ancient archaea (Lundquist *et al*., 2012b). Proliferation of homologs occurred in photosynthetic species and is likely linked to diversification in function of the ABC1K family due to the greater diversity of quinones present in photosynthetic species, including phylloquinone (vitamin K), ubiquinone, and plastoquinone (PQ9). Phosphorylation activity has not been shown directly for any ABC1K in bacteria or eukaryotes, although mutation of the human ABC1K homolog COQ8A (also named ADCK3) induced autophosphorylation *in vitro* (Stefely *et al*., 2015); it is conceivable that the ABC1K phosphorylate metabolites rather than proteins. PGs also contain metabolites important to photosynthesis, redox regulation, and defense against radical oxygen species (ROS) including phylloquinone, α-tocopherol (vitamin E), and plastochromonol-8 (PC8) as well as lipids and free fatty acids (van Wijk and Kessler, 2017). Although initially not considered part of the core PG proteome, lipoxygenase 3 (LOX3) involved in jasmonic acid biosynthesis, as well as pheophytin pheophorbide hydrolase (PPH) involved in chlorophyll degradation were found in PGs after stress treatment in the *abc1k abc1k3* loss-of-function mutant (Lundquist *et al*., 2013).

Genes showing similar mRNA expression patterns (from microarrays or RNAseq) across multiple conditions are often functionally related, *e*.*g*. acting in similar metabolic or signaling pathways or in complexes (Delli-Ponti *et al*., 2020; Rao and Dixon, 2019; Serin *et al*., 2016). However others have few significant co-expressers because they are regulated at a post-translational level. Construction of co-expression networks allows visualization of the correlation between the expression of genes and predict functions. Such networks can be constructed as ‘unforced’ networks if calculated from correlations of all genes within a dataset, or as ‘forced’ networks by choosing specific ‘bait’ genes of interest and pulling out their co-expressors and build edges from these selected baits and their co-expressors. Edges signify correlation between two nodes and are visually represented in a network by a line between two nodes. While ‘unforced’ networks are a good way to identify the strongest co-expression pattern(s) from a dataset, these correlations may cover up smaller co-expression relationships between genes of interest; a good example is the co-expression network for 10 geranylgeranyl diphosphate synthase paralogues (Ruiz-Sola *et al*., 2016) or organellar proteases (Majsec *et al*., 2017). Co-expression networks of both types have been widely used successfully, *e*.*g*. to discover core biotic stress responsive genes (Xie *et al*., 2011) (Amrine *et al*.), identify hub proteins in metabolic pathways and predict protein interactions (Ruiz-Sola *et al*.). Nearly 10 years ago, we generated a PG co-expression network for 25 proteins based on microarray data that helped understand PG functions (Lundquist *et al*., 2012a). The network was based on only 25 genes (baits) because there were no available mRNA expression data for the other PG proteins and because several PG proteins had not yet been identified. Currently 34 proteins have been assigned a PG localization (Table 1). Fortunately, far more complete co-expression data are now available through the most recent release of ATTED-II (https://atted.jp/) based on both microarray and RNAseq information (Obayashi *et al*., 2018). This provides a unique opportunity to revisit the PG co-expression network.

**Table 1.** The PG proteome composition with 34 proteins and selected features. Abbreviated names are used in the figures.

| **Table 1.** Proteins assigned to the PG proteome and used for protein and mRNA expression analyses |  |  |  |
| --- | --- | --- | --- |
| AT4G31390.1 | ABC1 kinase 1 (ABC1K1) | AT1G32220.1 | flavin-reductase related 1 (FDRED1) |
| AT1G79600.1 | ABC1 kinase 3 (ABC1K3) | AT4G38970.1 | fructose-bisphosphate aldolase-2 (SFBA-2) (d) |
| AT1G71810.1 | ABC1 kinase 5 (ABC1K5) | AT1G17420.1 | lipoxygenase 3 (LOX3) |
| AT3G24190.1 | ABC1 kinase 6 (ABC1K6) | AT1G78140.1 | methyltransferase related 1 (PGMT1) |
| AT3G07700.1 | ABC1 kinase 7 (ABC1K7) | AT2G41040.1 | methyltransferase related 2 (PGMT2) |
| AT5G05200.1 | ABC1 kinase 9 (ABC1K9) | AT5G13800.1 | pheophytin pheophorbide hydrolase (PPH) |
| AT1G06690.1 | aldo/keto reductase (AKRED) | AT1G54570.1 | phytyl ester synthase 1 (PES1) (b) |
| AT4G19170.1 | carotenoid cleavage dioxygenase (CCD4) | AT3G26840.1 | phytyl ester synthase 2 (PES2) (c) |
| AT5G41120.1 | esterase (ELT4) | AT3G27110.1 | plastoglobular M48 protease (PGM48) |
| AT4G04020.1 | fibrillin 1a (FBN1a) | AT3G10130.1 | SOUL heme-binding (HBP3) |
| AT4G22240.1 | fibrillin 1b (FBN1b) | AT4G32770.1 | tocopherol cyclase (VTE1) |
| AT2G35490.1 | fibrillin 2 (FBN2) | AT5G08740.1 | type II NADPH dehydrogenase (NDC1) (a) |
| AT3G23400.1 | fibrillin 4 (FBN4) | AT1G52590.1 | unknown protein (DUF393) |
| AT3G58010.1 | fibrillin 7a (FBN7a) | AT4G13200.1 | unknown protein 1 (U1) |
| AT2G42130.1 | fibrillin 7b (FBN7b) | AT3G43540.1 | unknown protein 2 (U2) (DUF1350) |
| AT2G46910.1 | fibrillin 8 (FBN8) | AT1G28150.1 | unknown protein 3 (U3) |
| AT2G34460.1 | flavin reductase-related 2 (FDRED2) | AT1G73750.1 | unknown SAG protein (SAG) |
| (a) dually targeted in plastid & mitochondria |  |  |  |
| (b) also Diacyl glycerol transferase 3 (DAGAT 3) |  |  |  |
| (c) also Diacyl glycerol transferase 4 (DAGAT 4) |  |  |  |
| (d) most of this protein is located in the stroma |  |  |  |

PG proteins were initially discovered from the analysis of PGs isolated from fully developed or senescent rosette leaf chloroplasts after plants were exposed to extended darkness, light stress or nitrogen deficiency (Lundquist *et al*., 2012a; Vidi *et al*., 2006; Ytterberg *et al*., 2006) and reviewed in (van Wijk and Kessler, 2017). Very little is known about the accumulation of these proteins in other tissue types, such as roots, embryos, various parts of the flower, siliques or seeds. However, a recent study determined relative protein abundance by mass spectrometry across 30 tissue types in Arabidopsis (Mergner *et al*., 2020) with data available via the ATHENA database (http://athena.proteomics.wzw.tum.de:5002/master_arabidopsisshiny/). As will be shown in this study, 33 out of 34 PG proteins were identified, providing a unique opportunity to address tissue type specific accumulation for PG proteins.

This study employs co-mRNA expression data from ATTED-II using both microarray and RNAseq information, as well as the proteomics data from ATHENA. Co-expressors were carefully annotated for subcellular location and function primarily based on the most recent literature and information in TAIR (https://www.arabidopsis.org/). Multiple co-expression networks were then generated by applying different co-expression thresholds to obtain biological insights for PG functions. Proteome abundance data from ATHENA were used for hierarchical clustering analysis to evaluate tissue-specific protein accumulation patterns. Combining this proteomics data with the mRNA co-expression networks and functional enrichment analysis provides new insights and research avenues into the different functions of PGs and its protein constituents.

## Materials and Methods

### Protein abundance and cluster analysis

To evaluate protein abundance for PG proteins across Arabidopsis tissues, relative protein abundance values (log2 iBAQ) were down-loaded from the ATHENA database (Mergner *et al*., 2020). Hierarchical cluster analysis for tissues and for proteins was then carried out for protein abundance across the tissue types after normalization by Zscore or Scaling (0-1). Significance scores for each cluster were calculated using the “pvclust” approach: https://github.com/shimo-lab/pvclust, providing “approximately unbiased” (AU) p-values, and clusters that have a high AU score (>= 95) can be judged to be strongly supported by the data. Cladograms were the same after normalization by Zscore or Scaling and there were only light differences in the AU p-values; only results using Zscore are shown.

### mRNA-based co-expression, networks and functional enrichment

Co-expressed genes for the PG proteins were downloaded (in the last week of July 2020) from the plant co-expression database ATTED-II (http://atted.jp/) (Obayashi *et al*., 2018) using the most recent dataset Ath-u1. This dataset is a unified version of co-expression calculated by linear regression of both RNA-seq and microarray co-expression data. The top-100 highest expressed genes based on the logit score (LS), which is a monotonic transformation of the Mutual Rank (MR) index, for each bait were used for detailed analysis. Larger LS indicates stronger co-expression, and LS=0 indicates no co-expression. Protein function was based on an updated version of the MapMan annotation system integrated into the PPDB. Protein experimental or predicted subcellular location was obtained from PPDB. Proteins were assigned to plastid, mitochondria, peroxisome or ‘other’. Updated information of function and/or subcellular location of the co-expressors was added to the PPDB http://ppdb.tc.cornell.edu/ (Sun *et al*., 2009). In case no annotation was predicted or experimentally determined, proteins were assigned to the location ‘other’. Functional enrichment tests for co-expressed genes for each bait and each network as a whole was based on the hypergeometric test (Majsec *et al*., 2017). Specifically, the relative functional distribution of the 35 functional bins was determined for each network and for the co-expressors of each of the PG baits within each network and compared to the predicted functions for the complete Arabidopsis proteome. Significant enrichment or underrepresentation was calculated using hypergeometric distribution. The minimal threshold was P < 0.05 with false discovery rate (FDR) multiple testing correction.

## Results and Discussion

### PG Protein abundance and distribution across 30 tissue types

The initial PG proteome reports for wild-type (col-0) Arabidopsis were published in 2006 (Vidi *et al*., 2006; Ytterberg *et al*., 2006) and updated in (Lundquist *et al*., 2012a). Proteome analysis of light stressed the *abc1k1 abc1k3* mutant show very strong PG enrichment of PPH and LOX3 (Lundquist *et al*., 2013; van Wijk and Kessler, 2017), bringing the total number of PG proteins considered here to 34 (Table 1). Importantly, these proteins were discovered from the analysis of PGs isolated from fully developed or senescent rosette leaf chloroplasts after plants were exposed to light stress. Very little is known about the accumulation of these proteins in other tissue types, such as root, various parts of the flower, siliques, seeds or embryos. A recent study determined relative protein abundance by mass spectrometry across 30 tissue types in Arabidopsis (Mergner *et al*., 2020) providing a unique opportunity to address tissue type specific accumulation for each of the PG proteins, with the exception of PES2 as it was not identified in this study. Hierarchical clustering to generate a cladogram for the tissue types based on protein abundance values (the few missing values were replaced by zero) and significance scores were calculated. Figure 1 shows these relative abundances as a heat map, with the proteins ordered by their average abundance and tissue types ordered following the cladogram. The cladogram showed a clear and consistent pattern, indicating that the underlying relative quantitative data were meaningful and that there is a coordinated accumulation of PG proteins across Arabidopsis parts and development. For example, the non-photosynthetic cell types (root, callus, cell cultures; 1-5) the stem tissues (hypocotyl, node and internode; 22-24), the various leaf tissues (petiole, cauline, shoot tip, lead distal and proximal; 15-20) each formed subclades. The non-photosynthetic samples of pollen (27), roots (1,9,10), callus (2,3), cell culture (4,5) and carpel (6) had the lowest overall average PG protein abundance (average and median values are listed in the bottom rows of the heat map), which supports the idea that PGs functions in particular in photosynthetic tissues as a conduit for thylakoid developmental transitions and facilitate metabolic flux into and out of the thylakoid membrane. LOX3, PPH, DUF393, SAG, lacked values for two or more tissues and these proteins had also on average some of the lowest average and median relative abundances (between 20 and 22). In particular PPH and SAG are known to be induced quite specifically during senescence, explaining why these were on average of low abundance. Furthermore, two additional proteins (ABC1K1 and U3) had no values for pollen. Several of the fibrillins (FBN1a, 1b, 2 and 4), together with UNKNOWN1 (U1) and SFBA2 had on average the highest relative abundance (between 27 and 30). A recent study confirmed PG-localization of U1 and a loss-of-function mutant showed impaired thylakoid biogenesis and reduced chlorophyll and carotenoids, but an increase in xanthophylls (Espinoza-Corral *et al*., 2019). Its molecular function remains to be determined. We note that most of the SFBA2 is located in the stroma with a minor, but persistent fraction co-purified with PGs (Lundquist *et al*., 2012a). Five proteins, PPH, SAG, PGM38, CCD4 and ABC1K7, are known to have the highest expression levels during senescence (van Wijk and Kessler, 2017) and are marked with arrows in Figure 1. CCD4 and PPH stand out because they showed the largest variation across sample types (calculated as cv) with highest values in senescent and cauline leaves, as well as silique valves, consistent with previous observations that their enzymatic function is most important under very specific conditions, namely senescence. Protease PGM48 showed the highest accumulation in the cotyledons, suggesting a particularly important role in this specialized organ. PGM48 was shown to be particularly important during leaf senescence with a postulated role in degradation of CCD4 (Bhuiyan *et al*., 2016; Bhuiyan and van Wijk, 2017). Pollen had the lowest average and median accumulation of PG proteins, but relative abundances of VTE1 and FRED1 were higher in pollen than all other tissue types, suggesting active prenyl lipid metabolism in pollen. We conclude that whereas PG proteins are present in most tissue types, they are most abundant in photosynthetic and senescing tissues. Subsets of PG proteins show higher accumulation levels in specific tissues types reflecting the different metabolic fluxes facilitated by PGs.

**Figure 1.**
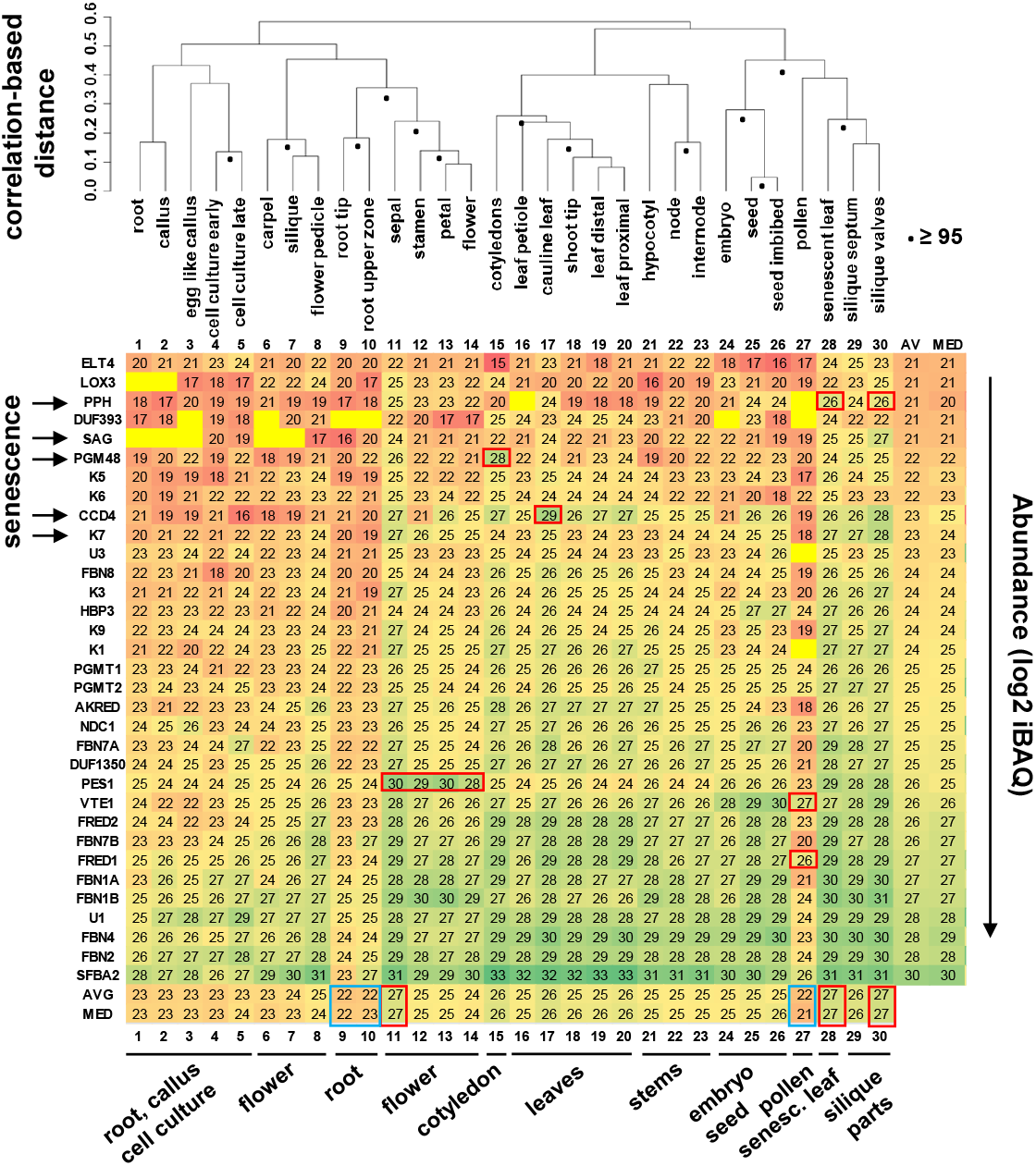
Heat map with relative abundance values of PG proteins listed in Table 1 (except for PES2 since it was not detected) across 30 tissue types in Arabidopsis from ATHENA (Mergner *et al*., 2020). Abundance values and protein annotations are listed in Supplemental Table 1. The abundance is calculated as log2 normalized iBAQ intensity. The iBAQ corresponds to the sum of all the peptide intensities divided by the number of observable peptides of a protein. Proteins in the heat map are ranked based on their average abundance across all tissue types, whereas the tissue types are organized based on hierarchical clustering after z-scoring of protein abundance vales within each tissue type. Missing values are marked up as bright yellow and lack of a value. Significance scores (AU) for each cluster were calculated and clusters that have a high AU score (>= 95) are indicated with a filled circle. Average (AVG) and median values (MED) are indicated. Color scale: bright yellow – missing value; low to high abundance from ref to green. Abundance value in this heat map range from 15 (darker red) to 33 (dark green). Tissue type numbering: 1 root; 2 callus; 3 egg like callus; 4 cell culture early; 5 cell culture late; 6 carpel; 7 silique; 8 flower pedicle; 9 root tip; 10 root upper zone; 11 sepal; 12 stamen; 13 petal; 14 flower; 15 cotyledons; 16 leaf petiole; 17 cauline leaf; 18 shoot tip; 19 leaf distal; 20 leaf proximal; 21 hypocotyl; 22 node; 23 internode; 24 embryo; 25 seed; 26 seed imbibed; 27 pollen; 28 senescent leaf; 29 silique septum; 30 silique valves.

### Co-expression networks and functional enrichment for the PG proteome

Previously we generated a top20 co-expression network for 25 PG proteins based on microarray data collected from MetaOmgraph (Singh *et al*., 2020) to help understand PG and plastoglobular protein functions (Lundquist *et al*., 2012a). The network was based on only 25 genes (baits) because at that time there were no mRNA expression data for the other PG proteins. Fortunately, far more complete co-expression data are now available through the most recent release of ATTED-II (http://atted.jp/) based on both microarray and RNAseq information. This provided a unique opportunity to revisit the PG co-expression network and re-evaluate enrichment for subcellular location and functions. We down-loaded 100 genes with the highest co-expression values (calculated as LS - see Methods) for each of the 34 PG genes (Supplemental Table S1A), resulting in a set of non-redundant 1553 genes (Supplemental Table S1B). We then constructed a co-expression network for the top20 highest co-expressors of each of the 34 PG genes (465 genes making 709 edges) to facilitate comparison to the top20 network that we published earlier for 25 genes (374 genes making 500 edges) (Lundquist *et al*., 2012a). We also generated co-expression networks based on three different minimal correlation thresholds for co-expression (LS ≥ 5, 6 or 7) with 987 (2061 edges), 564 (924 edges) and 282 (350 edges) non-redundant genes, respectively (Note that we limited the number of co-expressors to 100 for each PG gene). These threshold networks are likely more biologically meaningful than topN networks since they are controlled for significance of the co-expression, but dependent on the threshold can get very complex. The edges and non-reductant gene lists with associated information (including co-expression values and annotations for function and subcellular locations) for all four networks can be found in Supplemental Table S1A,B.

Figures 2, 3 and Supplemental Figure S2 show the annotated graphical displays for each of the four networks, clearly marking the PG genes with their abbreviated names, direct edges between PG genes (in red), and color coding for selected functions (colored edges) and subcellular location of co-expressors for plastidial, mitochondrial and peroxisomal proteins. Visual networks are most meaningful if baits and edges can be individually visualized, and we therefore show the LS ≥ 6 and LS ≥ 7 networks in Figure 3A,B. The LS ≥ 5 network is very dense due to high number of edges and is hard to visualize evaluate (Supplemental Figure S2). To better understand the direct interactions (edges) between the PG proteins themselves for LS ≥ 6 and 7, we visualized these direct edges at each threshold in a simple figure (Figure 4). This highlights that ABC1K3, ABC1K6, VTE1, AKRED, FDRED2 and PGMT1 are well connected to other PG proteins at LS ≥ 6. At LS ≥ 7, ABC1K1, ABC1K5, ABC1K6, VTE1, AKRED and FRED2 form a tight network (marked by the thicker, red edges), while ABC1K7 makes direct connections to ABC1K3 and PGM48. This suggested several sub-networks as indicated.

**Figure 2.**
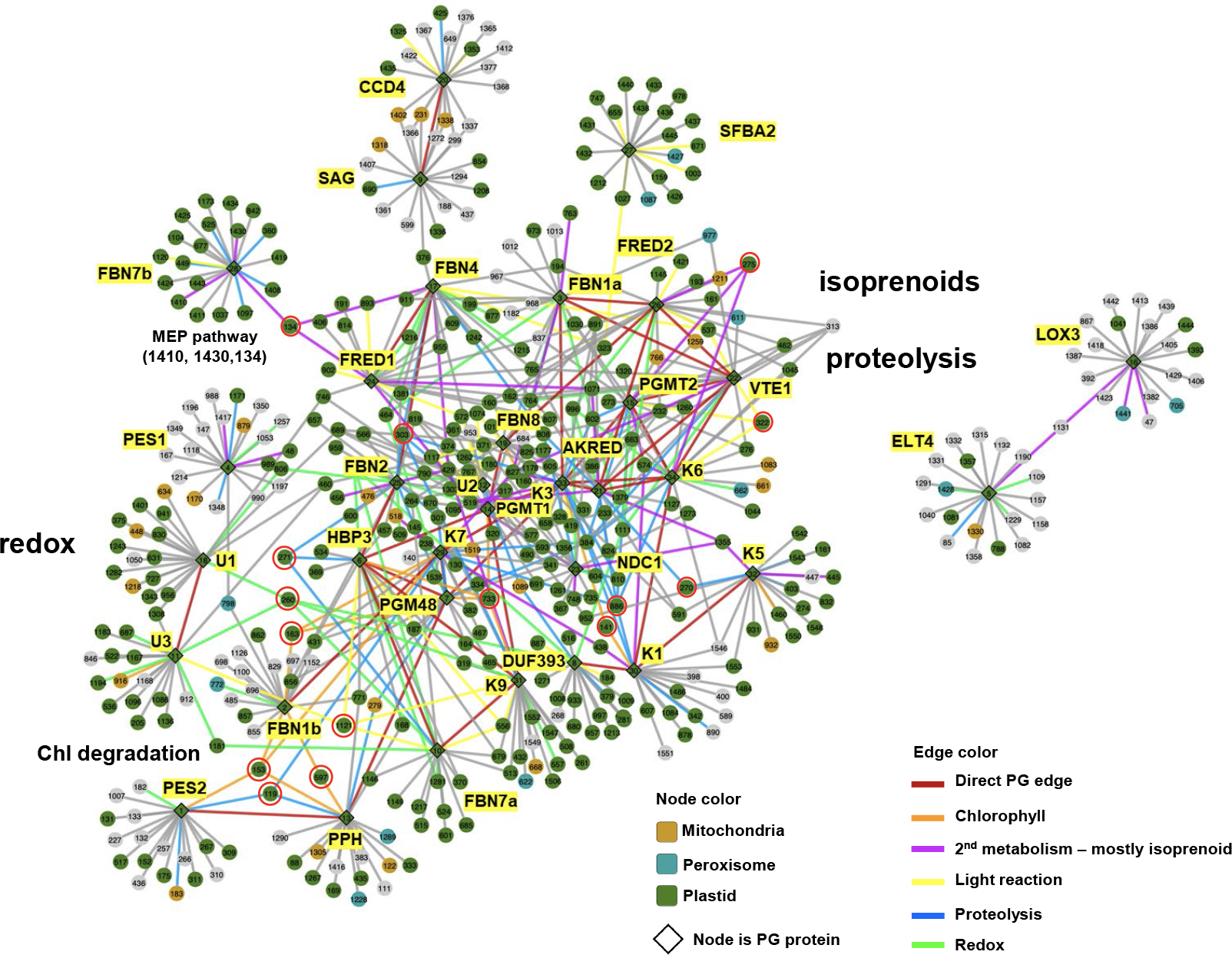
mRNA co-expression forced network for the PG proteome (34 proteins) based on combined micro-array and RNA-seq based gene expression data using the 20 genes with the highest correlation values (Logit Score Mutual Rank). Details for all co-expressors and correlations are listed in Supplemental Table S1A,B. Several co-expressors are marked by a red circle. Marked co-expressors that function in chlorophyll degradation are TIC55 (141), BCM2 (163), SGR1/NYE1 (153), PAO/ACD1 (597), PTC52 (733) – these show high connectivity to PG genes PES2, PPH, PGM48, ABC1K7 and FBN1b as well as the senescence-induced AAA+ ClpD chaperone (119). Co-expressors with a high number of edges (5 to 7) in this top20 network are: ferredoxin-thioredoxin reductase (260 - marked), glutaredoxin (896), HCF244 (891); thioredoxin-like/alkyl hydroperoxide reductase (1111), phytoene desaturase (PDS) (519), alpha/beta-Hydrolase (765), methionine sulfoxide reductase B1 (MSRB1) (232), FtsH1 (886-marked), FtsH2 (303-marked); FtsH8 (384). Other well visible co-expressors are: SPPA (270), LHCB7 (322), TAP38 (1121), MDS (134), SPS2 (275).

**Figure 3.**
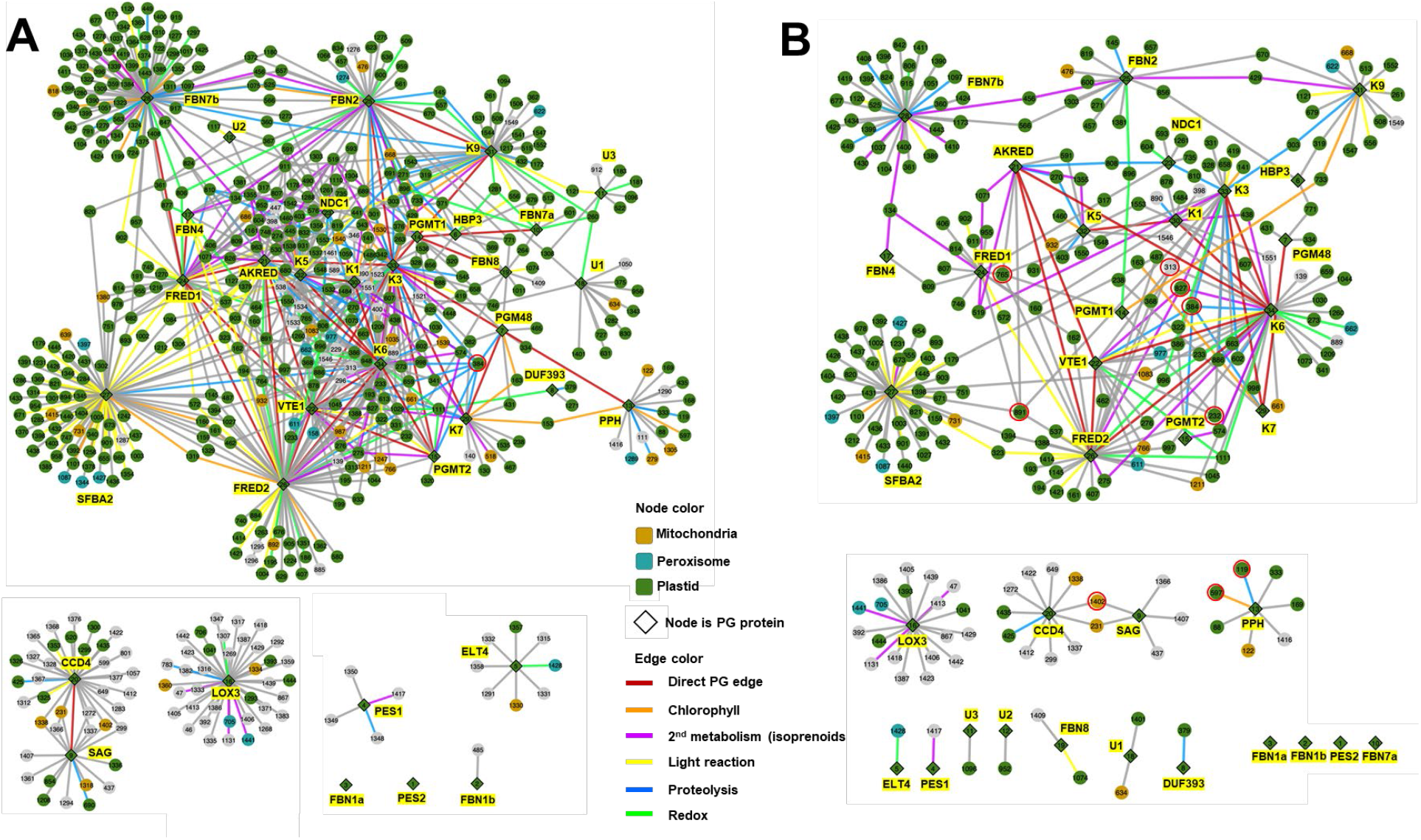
mRNA co-expression forced network for the PG proteome (34 proteins) based on combined micro-array and RNA-seq based gene expression data and minimal threshold correlation value (Logit Score Mutual Rank) of ≥ 6 (**A**) or ≥ 7 (**B**). Details for all co-expressors and correlations are listed in Supplemental Table S1A,B. Several co-expressors are marked by a red circle. In panel **A**, co-expressor FTSH8 (384) with 8 edges is marked. In panel **B**, co-expressors with the highest number of edges (4 or 5) are marked: FTSH8 (384), alpha/beta hydroxylase (765), HCF244 (891), methionine sulfoxide reductase B1 (232), squalene epoxidase (313) and ZEP (827). Other marked co-expressors are the MYBL2 Transcription factor (1402) involved in abiotic stress induced anthocyanin metabolism (with SAG and CCD4), chlorophyll degradation enzyme PAO (597) and senescence induced AAA+ chaperone ClpD with PPH. PG protein FDUF393 co-expresses tightly with the ClpS1 substrate adaptor (379) and ELT4 co-expresses with dehydroascorbate reductase-1 (1428).

**Figure 4.**
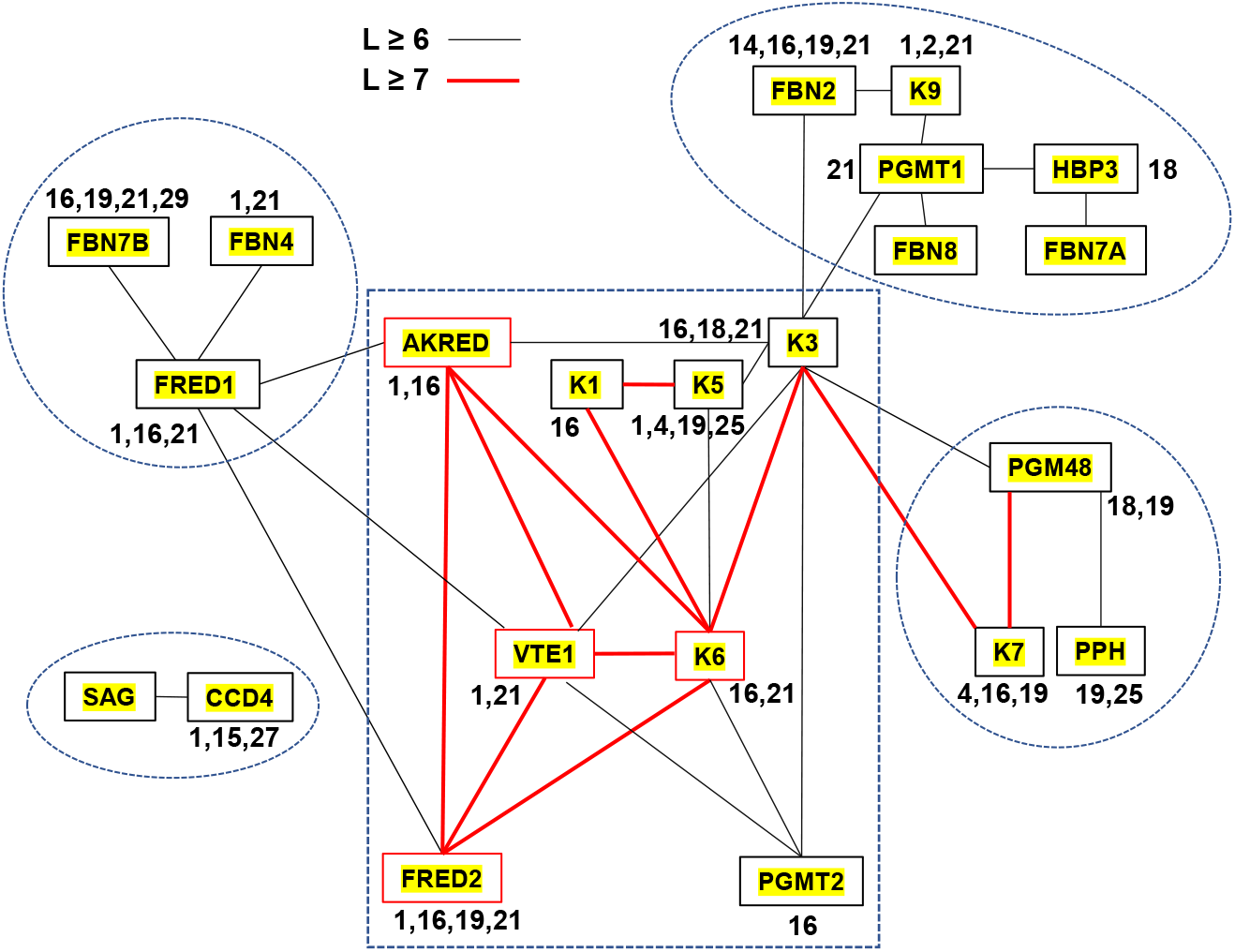
Diagram showing the direct co-expression edges between PG proteins based on the LS ≥ 6 and ≥7 networks. Red colored, thicker connections indicate edges that are also present in the more stringent network (L ≥ 7). Enriched functional bins are indicated for each PG protein. Explanations for Bin numbers are: 1 - Photosynthesis; 2 - Major CHO metabolism; 4 - glycolysis; 14 – S-assimilation; 15 - metal handling; 16 - secondary metabolism (mostly isoprenoids - bin 16.1); 18 - Co-factor and vitamin metabolism; 19 - tetrapyrrole synthesis; 21 - redox; 25 - C1- metabolism; 27 - RNA; 29 – proteostasis. The number of edges to other PG proteins at LS ≥ 6 / LS ≥7 are: K3 – 8/1; K6 – 6/4; VTE1- 6/3; FRED1 – 6/0; AKRED – 5/3; FDRED2 – 4/3; PGMT1 – 4/0; PGM48 – 3/1; PGMT2 – 3/0; K5 – 3/1; K7 – 2/2; K1 – 2/2; K9 – 2/0; FBN2 - 2/0; FBN4 – 1/0; FBN7A – 1/0; FBN7B – 1/0; PGM48 – 1/0; CCD4 - 1/0; SAG −1/0

We calculated enrichment for subcellular localization and function (using the MapMan bins; see PPDB) of the co-expressors for each of these networks and for the individual PG genes within each network (Table 2). All networks were highly enriched for plastid-localized proteins (66-79%), a small portion of mitochondrial proteins (5-6%) and peroxisomal proteins (2-4%), with the highest percentage of plastid proteins for the most stringent network (LS ≥ 7). Compared to the whole Arabidopsis proteome, in particularly photosynthesis (bin 1), 2^nd^ metabolism (bin 16; this was predominantly plastid isoprenoid metabolism - the MEP pathway, tocopherols, PQ9, PC8 and carotenoids), tetrapyrrole metabolism (bin 19; this was mostly chlorophyll degradation), and redox (bin 21) were enriched in the networks. In case of the LS ≥ 5 networks, enrichment was also observed for starch/sucrose metabolism (bin 2), glycolysis (bin 4), co-factors & vitamins (bin 18; mostly vitamin b6 (pyridoxine, pyridoxamine, pyridoxal) (Table 2).

**Table 2.** The number of co-expressors and enriched functions and subcellular localizations for the PG proteins in each of the four co-expression networks.

| Table 2. The number of co-expressors and enriched functions for the PG proteins at each of the co-expression networks |  |  |  |  |  |  |  |  |  |  |  |  |  |  |
| --- | --- | --- | --- | --- | --- | --- | --- | --- | --- | --- | --- | --- | --- | --- |
|  |  |  | # Co-expressors |  |  |  | Enriched functions* |  |  |  | subcellular local^ |  |  |  |
|  |  |  | top20 | LS ≥ 5 | LS ≥ 6 | LS ≥ 7 | top20 | LS ≥ 5 | LS ≥ 6 | LS ≥ 7 | top20 | LS ≥ 5 | LS ≥ 6 | LS ≥ 7 |
| Gene ID | ID# ** | whole network | 465 (681 edges) | 987 (2061 edges) | 564 (924 edges) | 282 (350 edges) | 1, 16, 18, 19, 21 | 1, 2, 4, 16, 18, 19, 21 | 1, 16, 19, 21 | 1, 16, 19, 21 | 69%; 5.6%; 2.8% | 66%; 6.1%; 2.0% | 75%; 5.7%; 2.5% | 79%; 5.0%; 3.5% |
| AT3G26840 | 1 | PES2 | 20 | 1 | 0 | 0 | 13, 19 | — | — | — | 50% | 0% | na | na |
| AT4G22240 | 2 | FBN1b | 20 | 10 | 1 | 0 | 19 | 19 | — | — | 45% | 50% | 0% | na |
| AT4G04020 | 3 | FBN1a | 20 | 9 | 0 | 0 | 16, 21 | 5 | — | — | 65% | 56% | na | na |
| AT1G54570 | 4 | PES1 | 20 | 17 | 4 | 1 | 4 | 4, 7, 8 | — | — | 15% | 18% | 0% | 0% |
| AT5G41120 | 5 | ELT4 | 20 | 24 | 7 | 1 | 21 | 21, 23 | — | — | 15% | 21% | 14% | 0% |
| AT3G10130 | 6 | HBP3 | 20 | 25 | 11 | 3 | 1, 21 | 21 | 18 | — | 100% | 96% | 100% | 100% |
| AT3G27110 | 7 | PGM48 | 20 | 34 | 11 | 4 | 18, 19, 21 | 18, 19 | 18, 19 | — | 95% | 94% | 100% | 100% |
| AT1G52590 | 8 | DUF393 | 20 | 22 | 3 | 1 | 1, 21 | 1, 21 | — | — | 90% | 86% | 100% | 100% |
| AT1G73750 | 9 | SAG | 20 | 40 | 17 | 5 | — | 18 | — | — | 30% | 43% | 29% | 0% |
| AT3G58010 | 10 | FBN7a | 20 | 30 | 8 | 0 | 1, 21 | 21 | — | — | 100% | 97% | 100% | na |
| AT1G28150 | 11 | U3 | 20 | 27 | 8 | 1 | 1, 4, 21 | 21 | 3, 4, 21 | — | 80% | 78% | 88% | 100% |
| AT3G43540 | 12 | U2 | 20 | 31 | 4 | 1 | 1, 4, 19 | 1, 29 | — | — | 90% | 94% | 100% | 100% |
| AT5G13800 | 13 | PPH | 20 | 50 | 15 | 7 | 5, 19, 25 | 5, 6, 19 | 19, 25 | — | 55% | 46% | 60% | 71% |
| AT1G78140 | 15 | PGMT1 | 20 | 49 | 17 | 4 | 21 | 18, 21 | 21 | — | 100% | 96% | 100% | 75% |
| AT2G41040 | 15 | PGMT2 | 20 | 51 | 13 | 3 | 16 | 16 | 16 | — | 80% | 75% | 85% | 100% |
| AT1G17420 | 16 | LOX3 | 20 | 70 | 38 | 19 | 16, 17 | 15, 16, 17 | 17 | 16, 17 | 15% | 14% | 13% | 16% |
| AT3G23400 | 17 | FBN4 | 20 | 52 | 10 | 2 | 1, 21 | 1, 21 | 1, 21 | — | 100% | 98% | 100% | 100% |
| AT4G13200 | 18 | U1 | 20 | 62 | 13 | 2 | 21 | 18, 21 | — | — | 80% | 81% | 31% | 50% |
| AT2G46910 | 19 | FBN8 | 20 | 57 | 9 | 2 | 1, 19 | 1, 19 | — | — | 90% | 95% | 89% | 50% |
| AT4G19170 | 20 | CCD4 | 20 | 71 | 31 | 9 | 1, 15 | 1, 15 | 1, 15, 27 | — | 25% | 31% | 29% | 22% |
| AT1G06690 | 21 | AKRED | 20 | 95 | 37 | 11 | 16 | 1 | 1 | 16 | 100% | 94% | 100% | 100% |
| AT4G32770 | 22 | VTE1 | 20 | 93 | 39 | 15 | 1 | 1, 16 | 1, 21 | 1 | 80% | 77% | 82% | 73% |
| AT5G08740 | 23 | NDC1 | 20 | 96 | 28 | 7 | 14, 21 | 14, 16, 29 | 14, 29 | — | 100% | 95% | 96% | 100% |
| AT1G32220 | 24 | FRED1 | 20 | 100 | 46 | 17 | 1, 16 | 1, 16, 21 | 1, 16, 21 | 1, 16 | 100% | 95% | 98% | 100% |
| AT2G35490 | 25 | FBN2 | 20 | 100 | 48 | 15 | 14, 21 | 14, 19, 21, 29 | 14, 19, 21 | 14, 16 | 95% | 95% | 94% | 93% |
| AT2G34460 | 26 | FRED2 | 20 | 100 | 78 | 34 | 1 | 1, 19, 21 | 1, 19, 21 | 1, 16 | 85% | 85% | 83% | 88% |
| AT4G38970 | 27 | SFBA2 | 20 | 99 | 79 | 40 | 1, 6 | 1, 8, 12, 19 | 1, 8, 19 | 1 | 90% | 88% | 91% | 98% |
| AT2G42130 | 28 | FBN7b | 20 | 99 | 85 | 33 | 16, 29 | 19, 21, 29 | 16, 19, 21, 29 | 16, 29 | 100% | 98% | 99% | 100% |
| AT3G07700 | 29 | ABC1K7 | 20 | 51 | 18 | 5 | 4, 16, 19 | 16, 18, 19 | 4, 16, 19 | — | 85% | 75% | 89% | 100% |
| AT4G31390 | 30 | ABC1K1 | 20 | 95 | 31 | 9 | 4, 19 | 2, 11, 16 | 16 | — | 70% | 67% | 68% | 78% |
| AT5G05200 | 31 | ABC1K9 | 20 | 100 | 41 | 12 | 1, 2, 21 | 1, 2, 18, 21 | 1, 2, 21 | 1 | 85% | 85% | 68% | 108% |
| AT1G71810 | 32 | ABC1K5 | 20 | 100 | 46 | 12 | 4, 19 | 1, 4, 25 | 1, 4, 25 | 19 | 85% | 73% | 70% | 83% |
| AT1G79600 | 33 | ABC1K3 | 20 | 100 | 54 | 23 | 19 | 16, 18, 21 | 16, 18, 21 | — | 95% | 82% | 93% | 96% |
| AT3G24190 | 34 | ABC1K6 | 20 | 100 | 68 | 40 | 16, 21 | 2, 16 | 16 | 16, 21 | 75% | 69% | 71% | 75% |

\* statistically enriched functions (only calculated if at least 10 co-expressors) using the MapMan bin numbers . 1 - Photosynthesis; 2 - Major CHO metabolism; 3 - minor CHO metabolism; 4 - glycolysis; 5 - fermentation; 6 - gluconeogenesis; 7 - OPP; 8 - TCA; 11 - lipid metabolism; 12 - N-metabolism; 13 - amino acid metabolism; 14 - S-assimilation (mostly Fe-S assembly); 15 - metal handling; 16 - secondary metabolism (mostly isoprenoids - bin 16.1); 17 - hormone metabolism (mostly Jasmonate); 18 - Co-factor and vitamins metabolism; 19 - tetrapyrrole synthesis; 21 - redox; 23 - nucleotide metabolism; 25 - C1-metabolism; 27 - RNA; 29 - protein
^ -subcellular localizations of co-expressors in the whole networks in the following order: plastid, mitochondria, peroxisomes. For each individual bait only plastid localization is shown.
\*\* identification number used in all networks

Table 2 also shows the number of co-expressors and their enriched functions, as well as plastid localization, for each PG protein in each network. The number of co-expressed genes at the least stringent co-expression threshold ranged from 1 to 100 (with 100 being the maximal number allowed). For example, PES2 has only one co-expressor at LS ≥ 5 suggesting that it is not part of a transcriptional network, whereas four of the ABC1Ks (ABC1K3,5,6,9), as well as FBN2, FBN7b, SFBA2 and the two flavin reductases (FRED1,2) had 99 or 100 co-expressed genes. With increasing stringency, the average number of co-expressors dropped, with some genes having no or just a few co-expressors but others still having more than 10 co-expressors (*e*.*g*. FBN6 and ABC1K6) at the highest threshold. The most frequently enriched functions for the individual PG genes were generally similar as for the whole networks. With increasing threshold stringency, the fraction of plastid-localized co-expressors increased for many PG genes – *e*.*g*. for ABC1K1 from 67% to 78%. However, PG proteins CCD4, ELT4, LOX3, PES1, PES2 and SAG did hardly co-express with plastid proteins; they have in common that they are most highly expressed during senescence (Table 2). The highest number of edges for co-expressors is 7, 8 and 5 for the top20, LS ≥ 6 and LS ≥ 7 respectively; this is consistent with highest overall connectivity of the LS ≥ 6 network compared to top20 and LS ≥ 7. The most connected co-expressors (that are not PG genes) are thylakoid protease FTSH8 (7, 8 and 5 edges), α,β- hydrolase AT4G36530 (7, 8 and 5 edges), and HCF244 involved in assembly of the Photosystem II reaction center (6, 8 and 5 edges) in the top20, LS ≥ 6 and LS ≥7 respectively.

In the following sections, we will use the graphical networks (Figures 2 and 3) and the direct PG edge network (Figure 4) to explore the role of individual PG proteins and their functional relationships to other PG proteins. We will focus on the top20 and LS ≥ 6, ≥ 7 networks totaling together 676 genes involved in 1134 non-redundant edges across these networks. Figure 4 also shows the statistically enriched functions of PG proteins for the LS ≥6, ≥7 networks.

### Comparison of the previous Top20 network with 25 PG genes and the new Top20 with 34 PG genes

In contrast to our previous network published in 2012 (Lundquist *et al*., 2012a), the new Top20 network shows that all PG genes, with the exception of ELT4 and LOX3, form a single network (Figure 2) (Supplemental Figure S1 for comparison of the two networks). For instance, VTE1 is now well connected to other PG proteins with direct edges, including ABC1K6, FRED2, PGMT2 and FBN1a, whereas in the previous top20 network VTE1 was not part of the PG network. Other general features of the prior top20 networks were conserved, such as direct edges between senescence induced CCD4 and SAG, but their connection was now much stronger with seven additional edges, as well as direct edges of PGM48 with PPH and ABC1K7 due to their (likely) role in chlorophyll degradation. The increased connectivity is also reflected in the higher number of edges per gene with 1.3 in the previous network and 1.5 in the new network. The improved connectivity likely reflects much richer and reliable mRNA expression data that is now assembled and utilized to generate co-expression data for Arabidopsis in ATTED-II. The network illustrates the enrichment of proteins involved in redox regulation (green edges) as well as proteins involved in isoprenoid metabolism (purple edges). Enrichment for members of proteolytic systems (blue edges) (in particular the thylakoid FTSH and stromal Clp systems) strongly overlays with the portion of the network enriched in proteins involved in isoprenoid metabolism. Enrichment for proteins involved in chlorophyll metabolism (in particular degradation) (orange edges) can be seen around PES2, PPH and FBN1b. Four PG proteins and ten co-expressors stand out for their high number of edges (between 5 and 7), all of them located in plastids. Importantly, four of these are PG proteins ABC1K1 (5x), ABC1K3 (6x), ABC1K6 (5x), VTE1 (5x) and together form the central hub of the PG network (Figure 4). Three of these high frequency co-expressors are members of the thylakoid FTSH protease complex (FtsH1,2,8), three are part of the plastid redox network (Fd-Trx-reductase A, glutaredoxin and a thioredoxin-like/alkyl hydroperoxide reductase), methionine sulfoxide reductase B1 involved in protein repair, phytoene desaturase (PDS) involved in carotenoid synthesis, HCF244, and a αβ-hydrolase with homology to PPH and chlorophyll dephytylase (CLD1) (see legend figure 2 for network id numbers). Several of these co-expressors are marked up in Figure 2 to illustrate their position in this network. Finally, we marked up 2 proteins (153 SGR1/NYE, 119 ClpD) associated with senescence and making edges between PES2, PPH and ABC1K7, and the thylakoid protein phosphatase (TAP38) involved in control of state transitions making edges to PG members FBN1b, FBN7a and HBP3 with demonstrated *in vitro* heme binding activity (Shanmugabalaji *et al*., 2020).

### Nuclear and plastid -transcriptional regulation and mRNA metabolism

Because the co-expression analysis clearly suggests coordination of the expression of subsets of genes, we first explored the presence of enriched transcriptional regulators and proteins involved in RNA metabolism (bin 27). Across the top20 and LS ≥ 6,7 networks there were 38 proteins in bin 27, of which 13 were plastid localized. Thirteen were found in all three networks, five of which were plastid proteins. This includes the well-studied Golden2-Like 1,2 nuclear transcription factors (GLK1,2) that are known to induce expression of genes involves in chloroplast biogenesis, including tetrapyrrole biosynthesis (Chen *et al*., 2016; Kobayashi and Masuda, 2016) and other nuclear TFs from different families (*e*.*g*. C2H2 zinc fingers and Basic Helix-Loop-Helix-bHLH). Plastid localized factors include several PPR and RRM proteins, as well as Sigma factor 5 (also named SigE) responsible for integrating light and circadian signals regulating chloroplast transcription (Belbin *et al*., 2017; Noordally *et al*., 2013), and the RNC1,3 RNA nucleases involved in mRNA processing (Hotto *et al*., 2015; Watkins *et al*., 2007). We then looked for transcriptional regulators as co-expressors that were shared between two or more PG baits as these are candidates to coordinate accumulation of subsets of PG proteins. Interestingly, the CCHC zinc finger protein AT5G20220 (#932 in the networks) co-expressed with ABC1K1 (LS = 6.0), ABC1K5 (LS = 7.7), ABC1K6 (LS = 6.4) as well as FRED2 (LS = 6.3). The function of this protein is not known but members of this family generally participate in RNA metabolism (Aceituno-Valenzuela *et al*., 2020). This protein (with multiple splice forms), as well as its single homologs in rice and maize, has a predicted mitochondrial location; however nearly all its co-expressors are plastid-localized, suggesting plastid location for one more splice forms (Supplemental Table S1). This suggests a link between plastid gene expression and expression of ABC1K1,5,6 and FRED2. The top3 co-expressors (LS ≥ 7) of this C2H2 zinc finger protein are thylakoid light harvesting protein LHCB7 (AT1G76570) expressed under very specific conditions (Klimmek *et al*., 2006; Peterson and Schultes, 2014), PG-localized ABC1K5 and plastid HCF145 (AT5G08720) involved in stabilization of the polycistronic psaA-psaB-rps14 (Manavski *et al*., 2015). The absence of LHCB7 is associated with lower rates of light-saturated photosynthesis and a diminished irradiance threshold for induction of photoprotective non-photochemical quenching; this very much aligns with a role of PGs in light stress response. Furthermore, zf-CCCH(5) protein AT3G02830 (#400) co-expressed (L ≥ 6) with ABC1K1 and ABC1K6, whereas plastid-localized PPR protein AT5G02830 (#1553) co-expressed tightly (LS ≥ 7) with both ABC1K1 and ABC1K5. Thus ABC1K1, ABCK5 and ABC1K6, but strikingly not the other three PG ABC1Ks (K3,7,9) are integrated with several plastid localized proteins involved in mRNA metabolism.

It was reported that ABC1K1 and ABC1K3 act oppositely in response to red light (Huang *et al*., 2015) (Yang *et al*., 2016). ABC1K1 was identified as a negative regulator downstream of Phytochrome B (PhyB) and bZIP nuclear transcription factor HY5 in red light mediated development, as evidenced by the suppression of the long hypocotyl elongation phenotype of *phyB* and *hy5* loss-of function mutants; ABCK3 reverses this effect of ABC1K1 (Yang *et al*., 2016). In other words, ABC1K promotes hypocotyl elongation in red light, whereas ABC1K3 inhibits this. The complex of COP1 E3 ligase and SPA (suppressor of phyA) homologs acts as a repressor of photomorphogenic responses by regulating the stability of photomorphogenesis-promoting transcription factors such as HY5, HFR1, LAF1 and CO, reviewed in (Podolec and Ulm, 2018). We therefore evaluated the co-expression networks for each of these factors. Interestingly, ABC1K1 (LS = 5.8) and ABC1K3 (LS = 6.3) co-expressed with SPA1 (AT2G46340; #1521), whereas SPA4 (At1g53090: #1368) co-expressed with ABC1K1 (LS = 7.6) and ABC1K6 (LS = 6.3). COP1 (At2g32950;1523) co-expressed with ABC1K1 (LS = 6.1). This does suggest that ABC1K1 and ABC1K3 and perhaps also ABC1K6 are part of a light signaling network. The co-expression networks did also include Phytochrome Interacting Factors (PIF) PIF4 (AT2G43010; #1337) and PIF5 (AT3G59060; #1328), but none of the other PIFs or light receptors (*e*.*g*. PhyA-E, CRY1,2, PHOT1,2, UVR8). PIF4, a key integrator of both light and temperature signaling pathways (Choi and Oh, 2016), was a co-expressor of both SAG and CCD4, whereas PIF5 co-expressed with just CCD4. MYB transcription factor AT1G71030 (#1402) co-expressed with both SAG (LS = 7.2) and CCD4 (LS = 8.1) consistent with these two PG proteins former a separate co-expression module at each of the three networks (top20, LS ≥ 6,7). We note that CCD4 co-expresses with some 10 transcriptional regulators (LS ≥ 6), including both Golden2-Like 1,2 (GLK1,2) nuclear transcription factors (#1376 and #1422) which are two partially redundant nuclear transcription factors that regulate chloroplast development in a cell-autonomous manner (Chen *et al*., 2016).

### The light and dark reactions of photosynthesis and photorespiration (bin 1)

Across the top20 and LS ≥ 6,7 networks there were 71 genes assigned to photosynthesis and photorespiration (bin 1) and involved in 103 edges. Table 2 shows which PG proteins are statistically enriched for co-expressors in bin 1. Overall, SFBA2 accounted to 45 of the edges within bin 1 (out of 103), whereas FRED1 and FRED2 accounted for 6 and 11 edges. We note that FRED1,2 also share a direct edge (LS = 6.3) (Figure 4) suggesting that they might act in concert. About 30% of the co-expressors in bin1 were involved with the stromal (dark) reactions, mostly the Calvin cycle, and they co-expressed nearly exclusively with SFBA2 and not with other PG proteins. SFBA2 also accounted for co-expression with many of the genes involved in the light reaction (26 edges), indicating that expression of the dark and light reactions is coordinated at the transcriptional level. As noted earlier, most SFBA2 is located in the stroma rather than associated with PGs. When excluding unique co-expressors to SFBA2, there were 5 co-expressors involved in the dark reactions and 28 involved in the light reactions distributed across the various complexes and functions (Photosystems, NDH complex, cyclic flow, state transitions and other regulators). Considering only the LS ≥ 7 network, FRED2 and ACB1K9 had most co-expressed genes (5 and 3, respectively) most of which involved in regulating of the light reactions (*e*.*g*. FLAP1, PIFI, TAP38) and cyclic electron flow components PGR5 and PGRL1A.

### Major carbohydrate metabolism - sucrose and starch metabolism (bin 2)

There were just 9 co-expressors in bin 2 making 11 edges across the three networks, but the pattern is striking and suggests that ABC1K9 plays a role in starch metabolism. The two major sucrose synthases (cytosolic) expressed in leaves (SPS1F/SPSA1 and SPS4F/SPSC) co-express with PES1 (SPSA1; LS = 5.1) or ABC1K5 and ABC1K6 (SPSC with LS = 7.5 and 6.2). These cytosolic enzymes are involved in degradation of maltose into sucrose and the double mutant for these synthases accumulates starch and high maltose indicating their critical role in starch degradation during the night (Volkert *et al*., 2014). Six of the nine genes are involved in starch metabolism and co-express with ABC1K9 (6≤ LS ≤7.3); since these genes do not make any edges to other PG proteins, this strongly suggests that ABC1K9 plays a role in connecting PG function to starch metabolism and warrants further investigation. Interestingly, starch synthase 4, co-expressing at high LS (7.3) with ABC1K9, physically interacts with FBN1a,b through its conserved N-terminus (Gamez-Arjona *et al*., 2014; Raynaud *et al*., 2016).

### Lipid and Fatty acid metabolism (bin 11)

There are 11 co-expressed genes involved in lipid metabolism making 16 edges, with two genes (FAD4 and HIL1) responsible for seven edges. FA desaturase 4 (FAD4) co-expresses with FRED2 (LS = 6.6), ABC1K6 (LS = 8.8) and VTE1 (LS = 6), whereas Heat Inducible Lipase 1 (HIL1) co-expresses with ABC1K3,6,7 (LS = 6.6, 7.7 and 6, respectively) as well as PGM48 (LS = 5.5). FAD4 is an unusual, substrate-specific palmitate desaturase introducing a delta-3 trans double bond at palmitate at the sn-2 position of phosphatidylglycerol, the only phospholipid in thylakoids. Although most of chloroplast PG assembly occurs at the inner envelope membrane, FAD4 is primarily associated with the thylakoid membranes facing the stroma (Gao *et al*., 2009; Horn *et al*., 2020). Peridoxin Q (PRXQ; AT3G26060) is required for this desaturation reaction, but it was not part of the co-expression networks (Horn *et al*., 2020). HIL1 is a plastid monogalactosyldiacylglycerol (MGDG) lipase that releases 18:3-FFA in the first committed step of 18:3/16:3-containing galactolipid turnover in particular during heat stress (Higashi *et al*., 2018). The strong co-expression with these four PG proteins supports the role of PGs in thylakoid lipid remobilization. Interestingly, PG-localized PPH (involved in chlorophyll degradation and phytol recycling) co-expresses with two peroxisomal proteins involved in β-oxidation (1289 - AT2G33150 and 1228 - AT4G29010), consistent with the role of PPH during senescence when lipids are metabolized.

### Isoprenoids (bin 16.1)

The three networks (top20 and LS ≥ 6, 7) include 26 genes involved in plastid-localized metabolism of different isoprenoids, making a total of 60 edges. Figure 5 summarizes plastid-localized isoprenoid metabolism and indicate proteins co-express with PG proteins. The figure includes the stromal MEP pathway that provides the C5 isoprene precursors (IPP and DPP), the central pathway for generation of C10, C15 and C20 isoprene diphosphates, pathways for metabolism of quinones, tocopherols, tocomoenols and tocotrienols, as well as carotenoids and xanthophylls. Finally, figure 5 (inset) also shows chlorophyl degradation leading to the release of phytol that can be used for new isoprenoid biosynthesis. This figure is mostly based on numerous recent reviews, in particular (Gutbrod *et al*., 2019; Wang and Grimm, 2021) (van Wijk and Kessler, 2017). The 26 co-expressors included six genes of the MEP pathway (CMK, MCT, DXR, HDR, HDS, MDS), eight genes involved in metabolism of tocopherols, tocotrienols, PQ9 or PC8, and 12 genes involved in carotenoid metabolism. A few genes generated four or more edges. These are PG protein VTE1 (6 edges), the MEP pathway genes MDS and HDS each with 4 edges, the carotenoid enzymes ZEP, ZDS and PDS with respectively 6, 5 and 5 edges, and SPS2 specifically involved in synthesis of PQ9/PC8 with 4 edges. 23 edges were formed with the six ABC1K PG proteins (ABC1K1 - 4x, ABC1K3 - 7x, ABC1K5 - 2x, ABCK6 - 6x, ABC1K7 - 3x and ABC11K9 - 1x), and other PG proteins with at least 4 edges were FBN7b (5x) and FRED1 (5x). Together, this clearly illustrate that PG proteins are heavily involved in isoprenoid metabolism forming a complex co-expression network. The challenge is to disentangle this network and arrive at more direct functional relationships between proteins and metabolites.

The MEP pathway enzymes are in particular co-expressed with the FBN family members FBN7b (4x), FBN2 (2x), and FBN4 (1x) perhaps because FBN proteins play a role in stabilizing isoprenoid metabolites. We note that stromal FBN5 is essential for PQ9 biosynthesis by binding to solanesyl diphosphate synthases (SPS) in Arabidopsis (Kim *et al*., 2015). The enzymes of the central pathway for generation of the isoprene diphosphates did not co-express with PG enzymes, perhaps since these are of general importance to a larger array of metabolites. The tocopherols, PQ9 and its downstream metabolites PC-8/mPC-8 share the common precursor homogentisate (HGA) and the enzymes HPPD, VTE1 and VTE3. Unique to each branch are SPS1,2 and HST for PQ9/PC8, and VTE2 for the tocopherols, tocomonoenols and tocotreinols (Figure 5).

**Figure 5.**
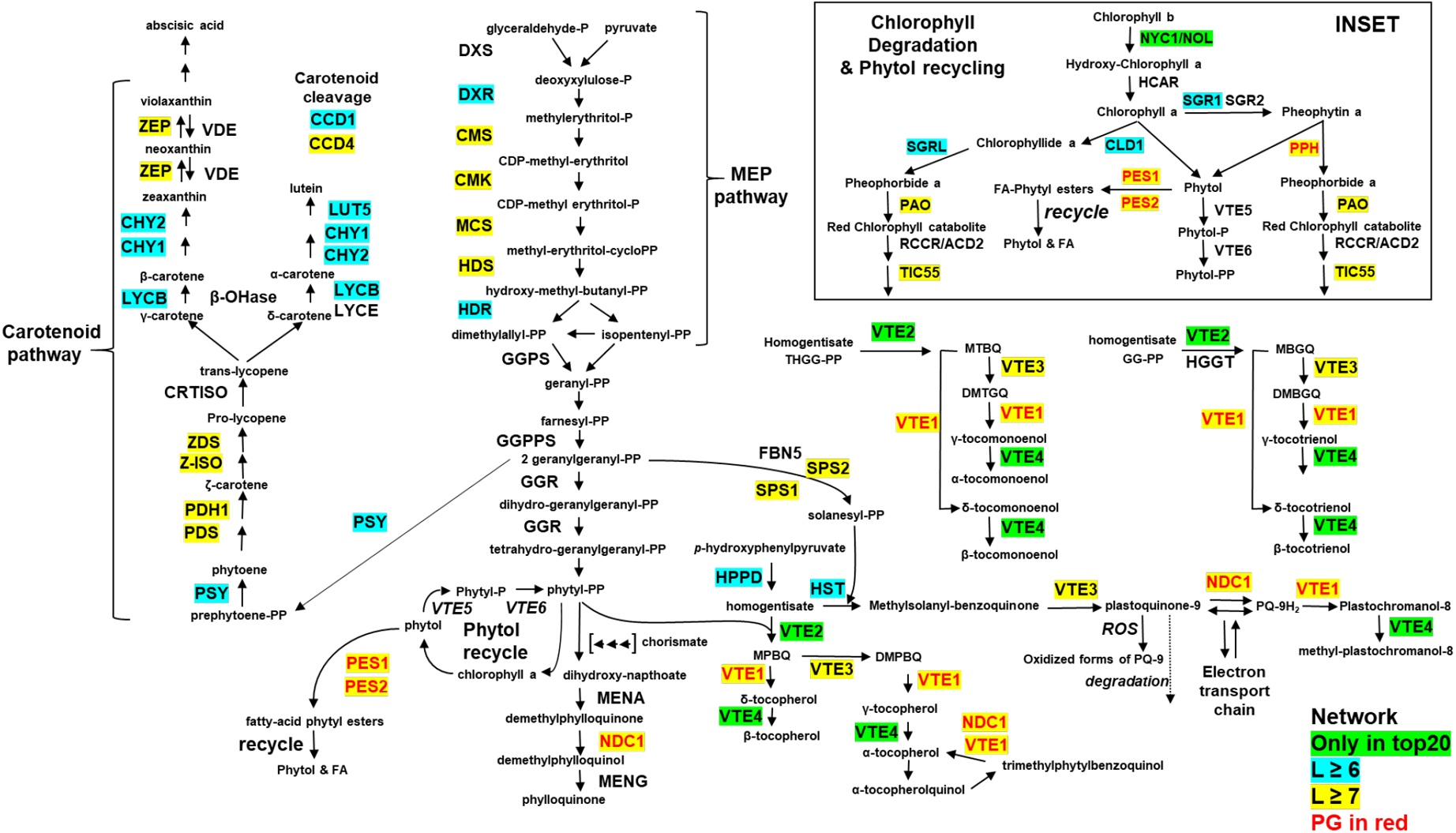
Overview of the metabolism of isoprenoid derived metabolites in the chloroplast. Plastid isoprenoid synthesis starts with the stromal MEP pathway that provides the C5 isoprene precursors (IPP and DPP). These C5 isoprenes are then condensed in a central pathway generating C10 (geranyl), C15 (farnesyl), C20 (geranyl-geranyl) diphosphates for the synthesis of quinones, tocopherols, tocomoenols and tocotrienols, as well as carotenoids and xanthophylls. The inset shows the plastid localized steps in chlorophyl degradation leading to the release of phytol that can be used for synthesis of phylloquinone and tocopherols. The major metabolites and enzymes are shown. Five of the enzymes are PG proteins (PPH, PES1, PES2, NDC1, VTE1) and they are shown in red letters. Proteins that are part of the three co-expression networks top20, LS ≥ 6 or LS ≥ 7 are marked up in green, blue or yellow, respectively.

Therefore, it is important to consider which of the PG proteins are co-expressors of these branch specific enzymes SPS1,2, HST and VTE2. SPS1 co-expressed with ABC1K1, ABC1K6 and PGMT2, SPS2 co-expressed with ABC1K6, PGMT2, FRED2, VTE1, whereas HST co-expressed with FRED1 and NDC1. VTE2 co-expressed only with one PG protein, namely PES1 and only in the top20 network at relatively low coefficient (LS = 5.4). VTE3 and VTE4 (involved in metabolism of most isoprenoids) only had one PG co-expressor each, namely SFBA2 for VTE3 and FBN1a for VTE4. This suggests that the function of the PG involves in particular in metabolism of PQ9/PC8 metabolism, and less directly in metabolism of tocopherols and tocotrienols. Most of the genes involved in carotenoid and xanthophyll biosynthesis co-expressed with PG proteins with high co-expression coefficients. Five of the PG localized ABC1K proteins (1,3,5,6,7), but not ABC1K9, accounted for 50% of all edges. This supports a tight coordination between PG functions and carotenoid metabolism as discussed previously (van Wijk and Kessler, 2017).

### Tetrapyrrole metabolism & chlorophyll degradation and recycling of phytol (bin 19)

The networks include 18 proteins involved in tetrapyrrole metabolism (Wang and Grimm, 2021), of which nine in synthesis and nine in degradation, forming a total of 29 edges. The inset of figure 5 shows the pathway of chlorophyll degradation, phytol esterification (by PES1 and PES2), and the recycling of phytol back to phytol-PP (by VTE5 and VTE6) for synthesis of tocopherols and phylloquinone. Five co-expressors are involved in the biosynthesis of protoporphyrin IX (GSA, GBP, UROM, CPO, PPO), upstream of the split between the heme and chlorophyl synthesis pathways. Thirteen proteins are exclusively involved with chlorophyll metabolism rather than heme metabolism (Supplemental Table S1). The five genes upstream of protoporphyrin IX, as well as Mg-protoporphyrin IX chelatase 2 (CHLI-2), all co-expressed with FBN7b; this striking result strongly suggests that this particular PG fibrillin is closely involved with early steps in tetrapyrrole biosynthesis. No other PG protein, except FNB2 with one edge to PPO, co-expressed with any of the biosynthetic enzymes upstream of protoporphyrin IX, reenforcing that FBN7b is somehow closely involved with these early biosynthesis steps. Chlorophyll degradation pathway proteins TIC55 (a hydroxylase) co-expressed with ABC1K1,3,6 as well as FBN2, SGR1/NYE1 co-expressed with PPH, ABC1K7, FBN1b and PES2, and PAO co-expressed with PPH and FBN1b, whereas PGM48 and PES2 both co-expressed with PPH. Chlorophyllide dephytalase CLD1 co-expressed with SFBA2, FBN8 and DUF1350. By far the tightest correlation was between PPH and PaO (LS = 10); these two proteins remove phototoxic metabolites in consecutive reactions likely explaining their tight co-expression. Indeed NYE, PPH and PAO have been shown to physically interact (forming a potential metabolom) (see (Aubry *et al*., 2020). Finally, A4G25650 with unknown function is a homolog of PAO and TIC55 and it co-expressed with six PG genes as PAO. This suggests that this PAO/TIC55 homolog is also involved in chlorophyll degradation.

### Vitamin biosynthesis (bin 18)

Five proteins involved in vitamin b6 synthesis (Asensi-Fabado and Munne-Bosch, 2010) were part of the three networks, together making 11 edges to 11 PG proteins, with an average correlation value of LS = 5.9. AT3G03890 with a Pyridox_oxidase domain stands out for having five edges with between 4.6 and 7.2 – PG proteins HBP3 and PGM48 had highest correlations. Vitamin b6 has plays a role in ROS defense, in particular as a quencher for singlet oxygen. The vitamin b6 deficient mutant *pdx1* have increased sensitivity to photooxidative stress and analysis of the *pdx1 vte1 npq1* mutant showed an interplay between tocopherols, carotenoids and vitamin b6 (Havaux *et al*., 2009). Pyridoxal 5’-phosphate (PLP) is an essential cofactor for glu-1-semialdehyde-amino transferase in the formation of ALA in tetrapyrrole synthesis and PLP is also a cofactor for starch phosphorylases and activates ADP-glucose-pyrophosphorylase (the first reaction in starch synthesis). Thus there are multiple potential explanations for the strong co-expression of the vitamin b6 pathways with PG proteome. Finally, phosphatase PYRP2 specifically involved in riboflavin (vitamin b2) biosynthesis (Sa *et al*., 2016) has three edges with ABC1K7, PGM48 and ABC1K3 (LS = 6.1, 6.2, 6.3); these three PG genes are also well-connected in the networks as is evident in figure 4.

### The chloroplast redox network and redox regulation (bin 21)

Thirty-two genes with a function in redox regulation are part of the co-expression networks forming a total of 67 edges. Most of these encoded for plastid localized proteins and four are located in peroxisomes. These genes include six plastid thioredoxins (f1, f2, m1, m2, m4, x), several glutaredoxins as well as components of the glutathione and ascorbate defense systems. In particular glutaredoxin AT5G03880 was highly connected with seven edges, gluthatione reductase with six edges and ferredoxin-thioredoxin reductase A, and an alkyl hydroperoxide reductase each with five edges across these three networks. At the highest stringency network (LS ≥ 7), there were 10 edges, with three from glutathione reductase connecting to FRED2, ABC1K3 and ABC1K6. This supports the concept that PGs are part of the redox and antioxidative network, likely able to accept and donate electron through the pool of PQ and other isoprenoids – for more discussion see (Havaux, 2020) (Pralon *et al*., 2020; Pralon *et al*., 2019) (van Wijk and Kessler, 2017) (Kumar *et al*., 2020).

### Protein turnover and proteostasis (bin 29)

There were some 112 genes assigned to bin 29 with a wide range in functions related to protein synthesis, folding, assembly, modifications and degradation. They formed a total of 236 edges involving all 34 PG genes. We focus on the highest stringency network (LS ≥ 7) to better understand the connectivity between the PG proteome and proteostasis functions. This narrowed it down to 51 genes (all but three encoding for plastid proteins) and 79 edges, distributed over six main functions, *i*.*e*. synthesis (29.1 & 29.2; 10x), sorting (29.3; 4x), PTM (bin 29.4 & 29.11; 7x), folding (bin 29.6; 4x), assembly (bin 29.8; 6x), processing and degradation (29.5; 20x). PG protein FBN7b had 14 edges, far more than any other PG protein, followed by ABC1K3 (7 edges), ABC1K6 (6 edges), FNB2 (5 edges) and NDC1 (5 edges). These co-expressors of FBN7b included five proteases (methionine aminopeptidase MAP1D, DEG8, Plsp1, ClpP4 and ClpP6), 4 ribosomal proteins and several involved in protein folding. The co-expressors with the most edges were HCF244 involved in D1 biogenesis (5x) making the three highest correlation edges to three PG reductases (AKRED, FRED1 and FRED2), thylakoid protease FTSH8 (5x) and protein repair protein methionine sulfoxide reductase B1 (MSRB1) (4x). HCF244 is a member of the atypical short-chain dehydrogenase/reductase superfamily and forms a complex with OHP1 and OHP2 likely aiding in delivery of pigments to the novo-synthesized D1 subunit of PSII (Hey and Grimm, 2020). OHP1 also tightly co-expresses with FRED2 (LS = 7.5), whereas OHP2 co-expresses with FRED2 and AKRED with LS = 5.9 and 5.1, respectively, supporting a functional connection between redox components of the PG and PSII biogenesis and repair. FTSH8 was very tightly correlated with ABC1K7 (LS = 10.5), followed by ABC1K6 (LS = 9.3) and ABC1K3 (LS = 8.2), and at lower correlations with VTE1 and UBI2 (LS = 7.7). Finally, within bin 29, two proteins stand out for their high correlation values namely the ATP-synthase assembly factor CGL160 (#513; LS = 9.2) (Fristedt *et al*., 2015) and plastid DWARF AND YELLOW 1 with unknown function (#319; LS = 10.8) (Huang *et al*., 2017) with shared edges to PG proteins HBP3 (LS = 7) and FBN2 (LS = 6.7).

### Lipoxygenase 3 and jasmonate metabolism and signaling

Jasmonate and its derivates are key regulators of senescence. Jasmonate synthesis starts in plastids through the action of lipoxygenase such as LOX3 (Wasternack and Kombrink, 2010). In contrast to most other PG genes, LOX3 co-expressors are mostly non-plastid proteins (only 13-16% plastid proteins; Table 2). LOX3 does not integrate into the main PG co-expression networks (Figures 2,3) but stands out for its four co-expressors with very high correlation values (LS ≥ 10). To put this in perspective, there are only 15 edges with correlation values ≥ 10 across the three networks, and four belong to LOX3. (Others are: ABC1K6, ABC1K7, ABC1K9, CCD4, FBN2, FBN7b, FRED1, FRED2, SFBA2 and PPH). Out of the 38 LOX3 co-expressors at LS ≥ 6 or 19 at LS ≥ 7, respectively 13 and 12 genes are in involved in jasmonate metabolism or signaling/responses (bin 17.2 and bin 17.7). These include the plastid localized JA synthesis enzymes LOX4, OAC1, OAC3 and AOS, peroxisomal 12-oxophytodienoate reductase (OPR3), as well as eight jasmonate-zim-domain proteins (JAZ1,2,5-10) involved in regulation of JA metabolism and response pathways. Perhaps explaining the recruitment of LOX3 (and other JA enzymes) to PGs is a recently proposed model where the chloroplast breakdown product pheophorbide a (generated by PPH – see Figure 5) acts a sensor for the rate of chlorophyll degradation and thereby regulates the speed of senescence tuned by JA signaling (Aubry *et al*., 2020).

## Conclusions

Taking advantage of the large accumulation of RNAseq data available now in the public domain, the ATTED-II database has generated mRNA co-expression data based on both RNAseq and microarray data and integrated this in a single co-expression coefficient (Obayashi *et al*., 2018). Based on this new information, we generated multiple co-expression networks for the PG proteome to better understand how the PG contributes to chloroplast homeostasis. The PG proteins formed a tight transcriptional network with ABC1K1,3, 5 and 6 together with VTE1 and two reductases (AKRED and FRED2) forming the center of the network. More peripheral to the network were a cluster with PGM48, PPH and ABC1K7 and a cluster with FRED1, FBN7B and FBN4. ELT4 and LOX3, as well as the pair SAG and CCD4 were disconnected from the main network. Several other PG proteins had no or very few co-expressors when applying a minimum threshold of co-expression (LS≥ 6 or 7); in such cases the top20 network was helpful to explore potential functional relationships.

A highly consistent functional enrichment was observed for the PG proteome regardless as to whether a top20 cutoff or a correlation threshold was applied to the co-expressors. Functional enrichment was observed in particular for chlorophyll degradation, isoprenoid metabolism, vitamin b6 metabolism, chloroplast redox regulation and proteostasis. Subsets of PG proteins associated with specific function, with in particular very strong relationships between chlorophyll degradation and PES2, ABC1K7, PGM48 and HBP3. Examples of surprising findings, providing working hypotheses for the study of molecular functions, were that ABC1K9 strongly co-expresses with many enzymes in starch metabolism, whereas FBN7b is involved with upstream steps of tetrapyrrole biosynthesis.

Most of the research on PGs has focused on chloroplasts from leaves in particular in response to various abiotic stresses (*e*.*g*. high light, low nitrogen), as well as senescence. However, PG have also been described for chromoplasts of pepper and tomato which suggested that chromoplast PGs mostly function to facilitate accumulation of carotenoids and contain high concentrations of biosynthetic enzymes whereas members of the fibrillin (FBN) family are involved in storing carotenoids (van Wijk and Kessler, 2017). The expression and protein accumulation of PG proteins in other non-photosynthetic parts or specialized parts such as cotyledons, parts of the influorescence, siliques, seeds and embryos have received very little research attention. The recent large scale proteome analysis of 30 different sample types from Arabidopsis (Mergner *et al*., 2020) provided an excellent opportunity to assess accumulation of PG proteins throughout the plant. Our analysis of relative protein abundance data of PG proteins (collected through the database ATHENA) showed that all PG proteins except PES2 were observed in most tissue types sampled and with clear differences in abundance across tissue types. Median and average value for the PG proteome were highest in green senescent tissues (leaves, sepals and silique valves). Several specific PG proteins showed a particularly high accumulation (unlike most of the other PG proteins) in selected tissues, such as the protease PGM48 in cotyledons, the carotenoid cleaving enzyme CCD4 in cauline leaves and VTE1 and FRED1 in pollen suggest specific targets in these tissues.

PGs act as microcompartments within plastids and in case of photosynthetic tissues, the lipid monolayer forms a continuum with the stromal-facing lipid layer of the thylakoid membranes. The PG-thylakoid connection thus allows exchange of lipophilic molecules (in particular isoprenoids, as well as fatty acids and membrane lipids), between the thylakoid and PG, whereas PG proteins have access to these small molecules. This allows bidirectional trafficking of small molecules such as phytol, PQ9, PC8, and tocopherols to adjust to the demands of the thylakoid electron transport chain, as demonstrated in the ABC1K1 and ABC1K3 mutants (Lundquist *et al*., 2013; Pralon *et al*., 2020; Pralon *et al*., 2019). Importantly, due to the high amount of PQ9 (and a lesser degree PC8 and tocopherols), the PGs also serve as a plastid oxidation-reduction buffer and redox exchange site, thus supporting ROS defense and oxidation-reduction reactions in metabolism (Havaux, 2020). Direct quenching of singlet oxygen by PQ9/PQH2 in thylakoids leads to oxidation products including PQOH9 and PQ(OH)39, and PGs serves as reserve pool and possibly a site of PQ repair and degradation (Ferretti *et al*., 2018; Ksas *et al*., 2015; Ksas *et al*., 2018). The PG could also serve as a site of detoxification of reactive carbonyls generated by ROS through the activity of oxidoreductases such as AKRED. Finally, the PG co-expression networks show the functional enrichment of the different thioredoxin systems that have shown to be so important in redox regulation for plastid metabolism, including tetrapyrrole biosynthesis (Wittmann *et al*., 2020).

Whereas the molecular functions of few PG proteins have been firmly established (*e*.*g*. VTE1, NDC1, PPH, PES1), the functions of most PG proteins remains enigmatic. The *in silico* analysis provided in this study has associated several PG proteins to specific metabolic processes and also provided a rich resource and foundation to experimentally elucidate the role of PGs in metabolism, plastid homeostasis and abiotic responses.

## Supplemental Data

**Supplemental Table S1**. Co-expressed genes and their functional annotation (from PPDB for 34 PG proteins in co-expression networks at different threshold levels from ATTED-II ath-u.1 for ATATH8.

**Supplemental Figure S1**. Side-by-side comparison on top20 mRNA co-expression forced networks for the PG proteome from this study and a previous study (Lundquist et al 2012)

**Supplemental Figure S2**. mRNA co-expression forced network (L≥5) for the extended PG proteome (34 baits) based on combined micro-array and RNA-seq based gene expression data and minimal threshold correlation value (ath-u.1 for ATATH8) from ATTED-II.

## Author contributions

EM generated the co-expression networks, carried out the functional enrichment analysis and contributed to the overall analysis and writing the paper. LP performed the hierarchical cluster analysis. KJVW conceived the study, annotated proteins for function and wrote the paper.

## Abbreviations

PGs: plastoglobules
LS: logit score (LS) which is the monotonic transformation of the Mutual Rank (MR) index
ROS: radical oxygen species
MS: mass spectrometry
MEP pathway: 2- C-methyl- d-erythritol 4-phosphate pathway;
PQ9: plastoquinone-9
PC8: plastochromonol-8

## Acknowledgements

We thank Dr Qi Sun for maintenance and support of the PPDB. We thank Kristina Majsec for advice on the construction of the co-expression networks and Cytoscape.

## Data availability statement

All essential data used for the proteome hierarchical clustering and co-expression are available in the Supplemental Data. Functional annotation is also freely available through the PPDB.

## Notes

### Competing Interest Statement

The authors have declared no competing interest.

